# Role of TRPV1 in colonic mucin production and gut microbiota profile

**DOI:** 10.1101/2020.04.17.046011

**Authors:** Vijay Kumar, Neha Mahajan, Pragyanshu Khare, Kanthi Kiran Kondepudi, Mahendra Bishnoi

## Abstract

**PURPOSE:** This study focuses on exploring the role of sensory cation channel Transient Receptor Potential channel subfamily Vanilloid 1 (TRPV1) in gut health, specifically mucus secretion and microflora profile in gut.

**METHODS AND RESULTS:** We employed resiniferatoxin (ultrapotent TRPV1 agonist) induced chemo-denervation model in rats and studied the effects of TRPV1 ablation on gut mucus secretion patterns. Histological and transcriptional analysis showed substantial decrease in mucus production as well as in expression of genes involved in goblet cells differentiation, mucin production and glycosylation. 16S metagenome analysis revealed changes in abundance of various gut bacteria, including decrease in beneficial bacteria like *Lactobacillus spp* and *Clostridia spp.* Also, TRPV1 ablation significantly decreased the levels of short chain fatty acids, *i.e.* acetate and butyrate.

**CONCLUSION:** The present study provides first evidence that systemic TRPV1 ablation leads to impairment in mucus secretion and causes dysbiosis in gut. Further, it suggests to address mucin production and gut microbiota related adverse effects during the development of TRPV1 antagonism/ablation-based therapeutic and preventive strategies.

## 1. INTRODUCTION

Transient receptor potential channel vanilloid 1 (TRPV1) is a non-selective cation channel, that transports mostly Ca^2+^, also Mg^2+^ and Na^+^. It is activated by a variety of stimuli, such as heat (<40°C), acidic pH and also various dietary components. Capsaicin from chilli, has been studied a lot in relation to its beneficial effects in metabolic complications like obesity. In most studies, capsaicin is shown to improve the composition of gut microflora towards health-promoting bacteria (Baboota et al., 2014; Baskaran et al., 2016; Zheng et al., 2017; Zsombok and Derbenev, 2016).

In gut dysbiosis during obesity, colitis etc, pathogen-initiated depletion of mucus layers and infection is observed, resulted from a decrease in beneficial bacteria and increase in population of harmful pathogenic bacteria. (Ng et al., 2013; Pacheco et al., 2012; Png et al., 2010). Such conditions were seen to be improved with capsaicin administration. Recently, capsaicin was shown to upregulate MUC2 gene in the intestine, with an increase in population of mucin-feeding beneficial bacterium *Akkermansia muciniphila* (Baboota et al., 2014; Shen et al., 2017). Further, there are studies relating capsaicin to mucus secretion in respiratory tract, and role of TRPV1 in increased expression of mucin genes MUC2, MUC5AC was observed (Yang et al., 2013). In other literature, mucus secretion has been shown to be regulated by inflammatory cytokines, neurotransmitters and hormones (Plaisancie et al., 1998); and interestingly, many of these molecules are also known to be affected by TRPV1 modulation. TRPV1 positive neurons profusely innervate the gut, including intestines, and mediate responses to stimuli via local release of neurotransmitters and/or by communicating to the central nervous system (Holzer, 2008; Plaisancie et al., 1998).

TRPV1-knockout models have been extensively used in cancer and inflammation research (Bode et al., 2009; Bujak et al., 2019; Fernandes et al., 2012; Santoni et al., 2012; Toledo-Maurino et al., 2018). Alternatively, high doses of capsaicin or resiniferatoxin (RTX), an ultrapotent agonist of TRPV1, have been employed to achieve systemic TRPV1 denervation and study its effect on pain and inflammation, chronic pain management in diseases like cancer (Bujak et al., 2019; Fukushima et al., 2017; Jeffry et al., 2009; Mishra and Hoon, 2010; Pecze et al., 2017). But there has been limited focus on gut health and metabolism studies in such models. No research has explored how TRPV1 denervation might affect mucus secretion in gastrointestinal tract.

Based on the available literature, we hypothesized that TRPV1 may have an active role in maintenance of mucus secretion in gut, directly or indirectly, which further affects the overall health and metabolism of an individual. In present study, we have employed systemic TRPV1 chemo-denervation model in rats using RTX, to explore the effect of TRPV1 ablation from body, on mucus secretion patterns in gut. We examined the changes in gut mucus secretion and related parameters in absence of TRPV1^+^ neurons, and explored its effect on gut microbiota and metabolism. Here, we provide first evidence that TRPV1 denervation negatively affects the parameters of mucus production in gut and disturbs the gut bacterial profile.

## 2. MATERIAL AND METHODS

### 2.1. Animals

Six weeks old male Wistar rats were procured from IMTech Center for Animal Resources and Experimentation (iCARE), Chandigarh, India, and housed in Animal Experimentation facility at National Agri-Food Biotechnology Institute (NABI), Mohali, India. Animals were kept in pathogen-free environment at 25±2°C, and maintained on a 12-hour light-dark cycle. They were divided into two groups – Control and RTX (Resiniferatoxin~95%; Sigma-Aldrich, Missouri, United States) (n=3 each). All animals were given free access to water and normal pellet diet throughout the experiment. Based on our power analysis (previous studies and other available literature on RTX induced chemo-denervation, n=3 is sufficiently appropriate (100% animals are showing denervation) number for our experiments. Experimental protocol was approved (Approval number NABI/2039/CPCSEA/IAEC/2019/04) by Institutional Animal Ethics Committee (IAEC) of NABI. Experiment was conducted according to the Committee for the Purpose of Control and Supervision on Experiments on Animals (CPCSEA) guidelines on the use and care of experimental animals. Plan of experiment is briefly explained in Figure 1.

**Figure 1.**
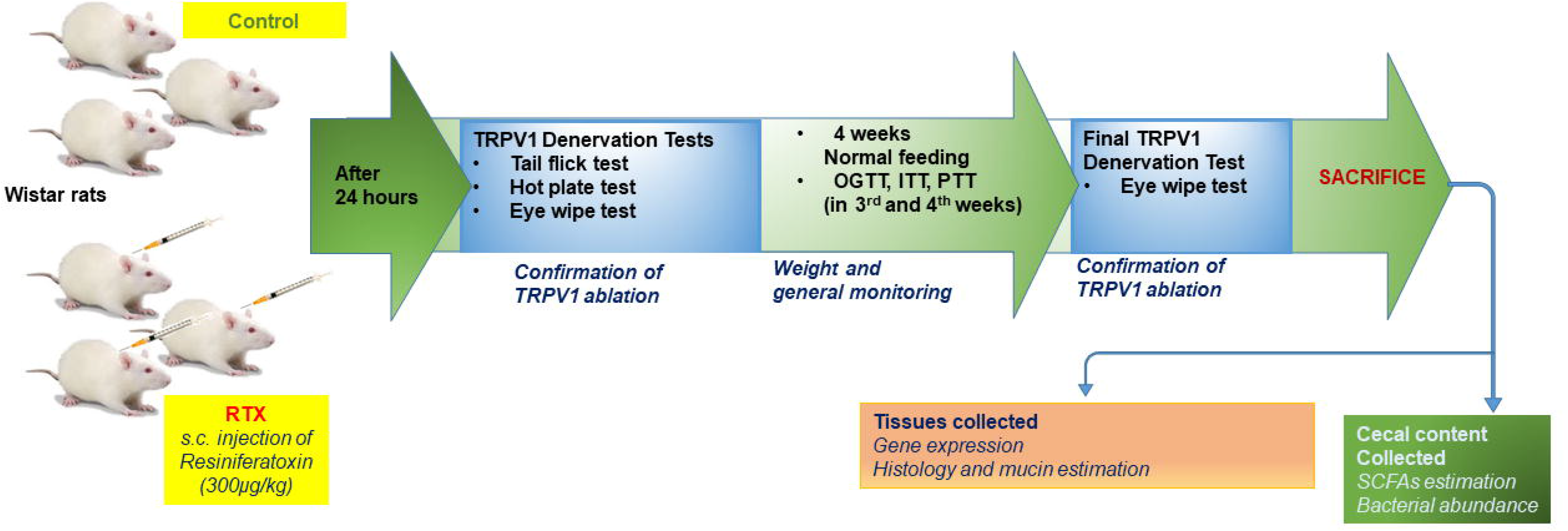
General experimental plan.

### 2.2. Chemo-denervation and confirmation tests

Systemic ablation of TRPV1^+^ neurons in RTX group animals was achieved by single subcutaneous injection of maximum tolerable dose of RTX; 300μg/kg body weight of animal (Szallasi and Blumberg, 1992; Szallasi et al., 1989), which was confirmed by loss of physiological responses after 24 hours. Various tests were employed for confirmation – Tail flick test, Hot plate test, Eye wipe test. For tail flick test, dolorimeter with 2-3A current was used and time of tail flick due to heat was noted. Cut-off time was set at 12 seconds. In hot plate test, the surface temperature was kept at 55±5°C and number of jumps/paw licks were recorded for 15 seconds. Experiments were repeated 3 times per animal, at intervals of 5 minutes. For eye-wipe test, 0.02% w/v capsaicin solution (capsaicin≥95%; Sigma-Aldrich, Missouri, United States) was used and number of wipes was recorded for 30 seconds. Replicates for each animal were taken using both eyes separately.

### 2.3. Blood glucose measurement

Oral glucose tolerance test (OGTT), insulin tolerance test (ITT) and pyruvate tolerance test (PTT) were done during 3^rd^ and 4^th^ weeks of study for measurement of blood glucose and assessment of glucose homeostasis. Doses used for tests were 2g/kg body weight p.o. glucose, 1U/kg i.p. insulin and 1g/kg i.p. sodium pyruvate respectively. Blood glucose levels were measured at 0 (before treatment), 15 min, 30 min, 60 min, 90 min and 120 min after treatment, using Glucocard (Arkray, Japan).

### 2.4. Histological analysis

Upon sacrifice, colon and ileum tissues were stored in 10% formalin for further processing. For paraffin embedding, tissues were prepared by serial dehydration in ethanol (25%, 50%, 70%, 90%, 100%, 2 hours each) followed by xylene treatment (2hours, twice). After embedding (in molten paraffin at 60°C, 2 hours, twice) and microtomy blocks formation, 5μm thick sections were made. For staining, sections were de-paraffinized using xylene (2-5 min, twice) and subjected to serial rehydration in ethanol (100% to 25%, 2 min each, followed into water). Alcian blue-Hematoxylin-Eosin (Hi-media Laboratories, Mumbai, India) stains were used for 15 min, 30 sec, 15 sec respectively, with washing (thrice in water for 5 min each, after every staining step). Slides were then again serially dehydrated and treated in xylene, and mounted using DPX mountant (Hi-media Lab, Mumbai, India). Tissue morphology and mucin production were observed. The analysis for Alcian blue intensity (mucins staining) was done using software ImageJ (from https://imagej.nih.gov/).

### 2.5. Gene expression analysis

RT-qPCR was employed for gene expression analysis in colon, ileum and dorsal root ganglia samples. Total RNA was isolated from tissues using Trizol-Chloroform-Isopropyl alcohol method and quantified on Nanodrop (Thermo Fisher Scientific, Massachusetts, United States). RNA integrity was determined by agarose (1.2%) gel electrophoresis and DNase (Thermo Fisher Sci.) treatment was given to eliminate any genomic DNA contamination. cDNA synthesis was done using RevertAid First Strand cDNA Synthesis Kit (Thermo Fisher Sci.) and relative change in gene expression was determined by qPCR using SsoAdvanced Universal SYBR Green Supermix (Bio-Rad, California, United States). qPCR was performed on CFX96 Touch Real-Time PCR Detection System (Bio-Rad) under following conditions: initial denaturation – 95°C, 2 min, [denaturation – 95°C, 5 sec; annealing/extension –60°C, 30 sec] x 40 cycles, final extension – 60°C, 5 min and melt curve analysis between 60°C-95°C with 0.5°C/5 sec increment. Data was analyzed using ΔΔCt method (Livak and Schmittgen, 2001), β-actin, GAPDH or Ubiquitin C (Ubc) genes were used for normalization. The list of primers used in the experiment is given in Table 1.

**Table 1:**
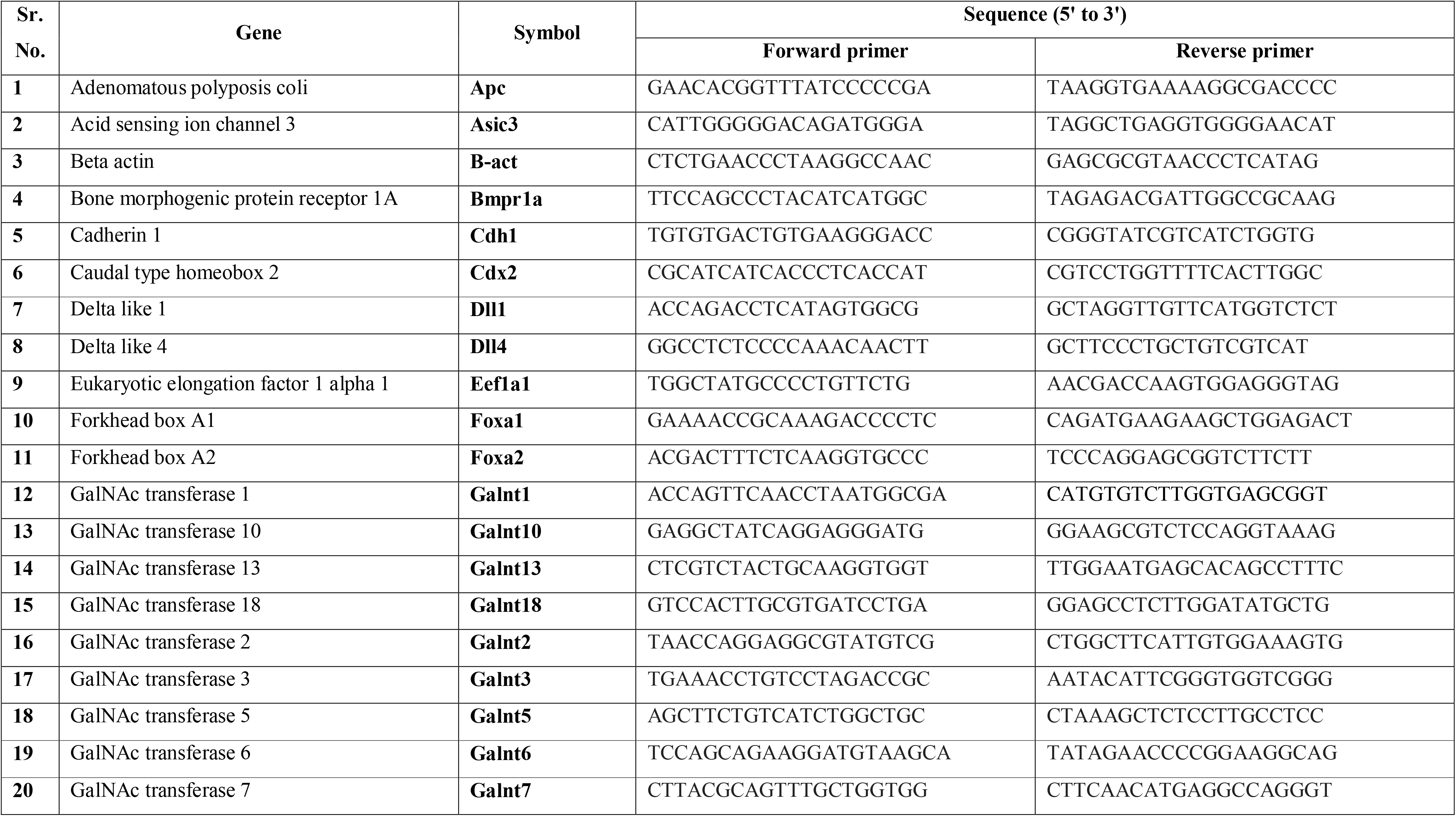
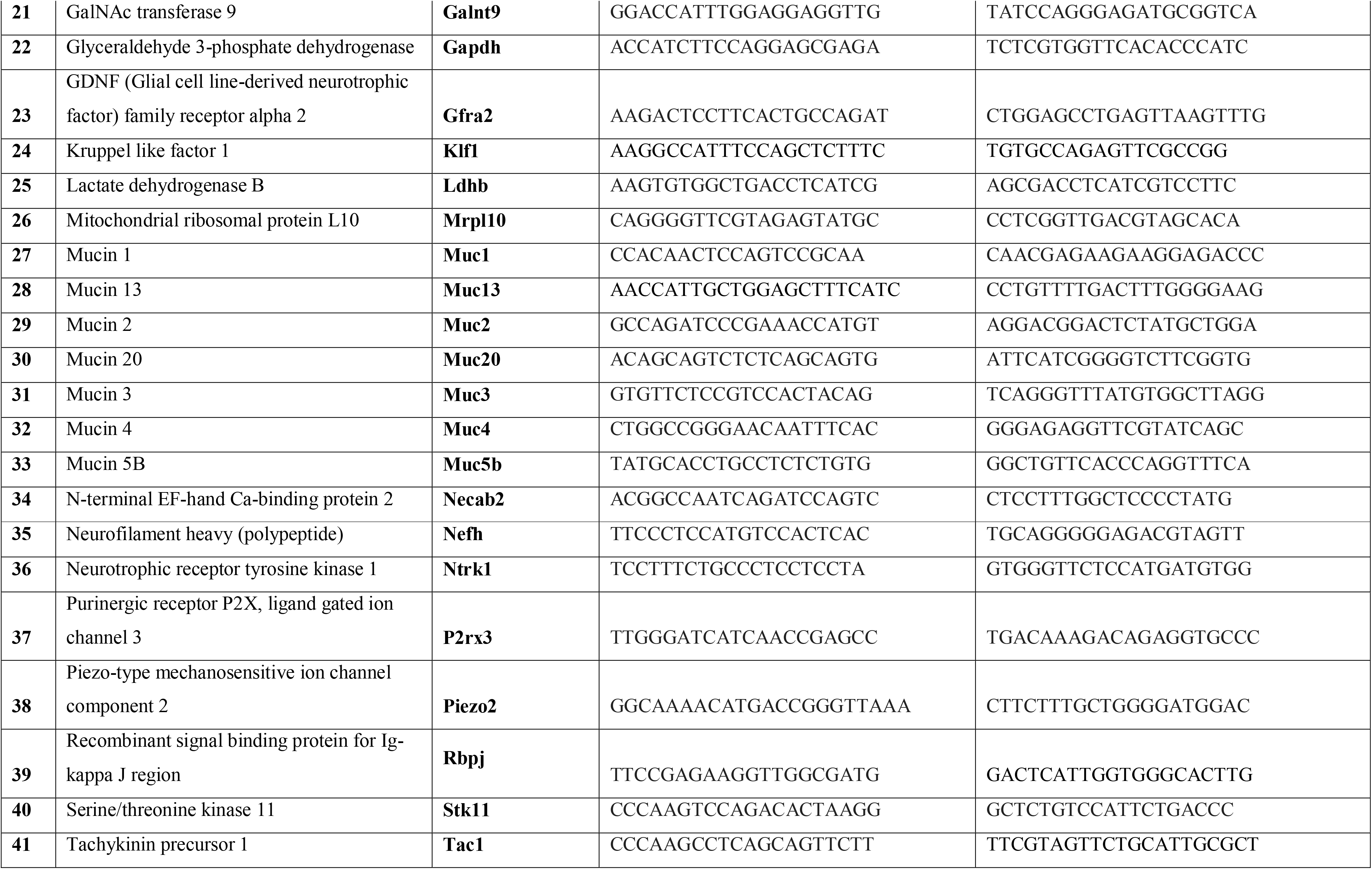
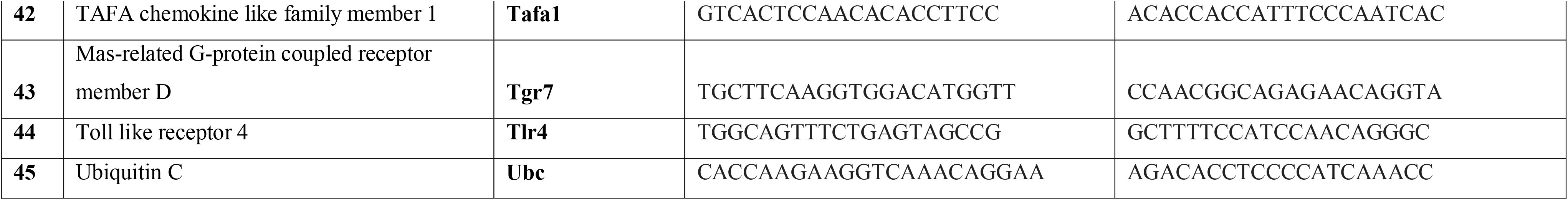
List of primers used in the study.

### 2.6. Short chain fatty acids (SCFAs) estimation

SCFAs were measured in cecal content of rats using HPLC. Agilent 1260 Infinity series chromatographic system (Agilent Technologies, Singapore) with previously described method (Singh et al., 2018) was employed. In short, 100-120 mg samples were homogenized in 500μl acidified water each (pH 2) by thoroughly vortexing, kept at room temperature for 10 min and centrifuged at 6000 rpm, 20 min, 4 °C. Supernatant was filtered using Millex-GN 0.2 μm nylon syringe filters (Millipore, Massachusetts, United States). 20 μL samples were injected into Hi-Plex H column (300×7.7 mm; 8 μm particle size, Agilent Tech.). 0.1% formic acid in Milli-Q water (Merck Millipore, 0.22 μm filtered, resistivity 18.1–18.3 MΩ cm) was used as mobile phase. The column was equilibrated and eluted with an isocratic flow rate of 0.6 mL/min at 50 °C for 60 min. Volatile Acids Mix (Cayman Chemicals Co., Michigan, United States) was used as standard at concentration range 100-6400μM. Final sample concentrations were calculated as μM/mg.

### 2.7. Bacterial abundance

16S metagenome analysis was done in cecal content. The bacterial genomic DNA was isolated from cecal content of rats using NucleoSpin DNA Stool kit (Macherey Nagel, Düren, Germany), following the manufacturer’s instructions, which was used in 16S metagenome analysis on Illumina sequencing platform (outsourced to NGB Diagnostics Pvt. Ltd., New Delhi, India). Analysis was done using online available resources (https://rdp.cme.msu.edu, www.eztaxon.org).

### 2.8. Statistical analysis

Data was analyzed using Graphpad Prism software (Graphpad, San Diego, California, USA). All data is presented as mean±SEM (bar graphs, line graphs) or individual values (dot plots). Student’s t-test (unpaired, with Welch’s correction) was employed to assess the differences between the two groups. P<0.05 was considered significant in all data (however, numerical value is also given at certain places to include some major but statistically non-significant changes).

## 3. RESULTS

### 3.1. Selective TRPV1 ablation disrupted normal glucose homeostasis

RTX caused total loss of physiological responses to noxious heat or capsaicin in rats. In tail flick test, no response to heat was observed in RTX group throughout test time, compared to very short latency times shown by control group animals (Figure 2A). Similarly, in hot plate test, there was no response (jumps or paw licks) from RTX group animals, indicating complete loss of sensitivity to temperature of 55±5°C (Figure 2B). These tests proved the systemic TRPV1 ablation in treated animals. To see the response to capsaicin-induced noxiousness/pain, 0.02% w/v capsaicin solution was used in eye wipe test. Following same pattern as before, there was no response at all, from the RTX group animals, compared to frequent eye wipes by control rats (Figure 2C). These tests were in agreement with the literature regarding TRPV1 denervation by RTX. Also, the eye-wipe test showed same results even after 4 weeks (Figure 2D), confirming that there was no or negligible recovery of TRPV1^+^ neurons.

**Figure 2.**
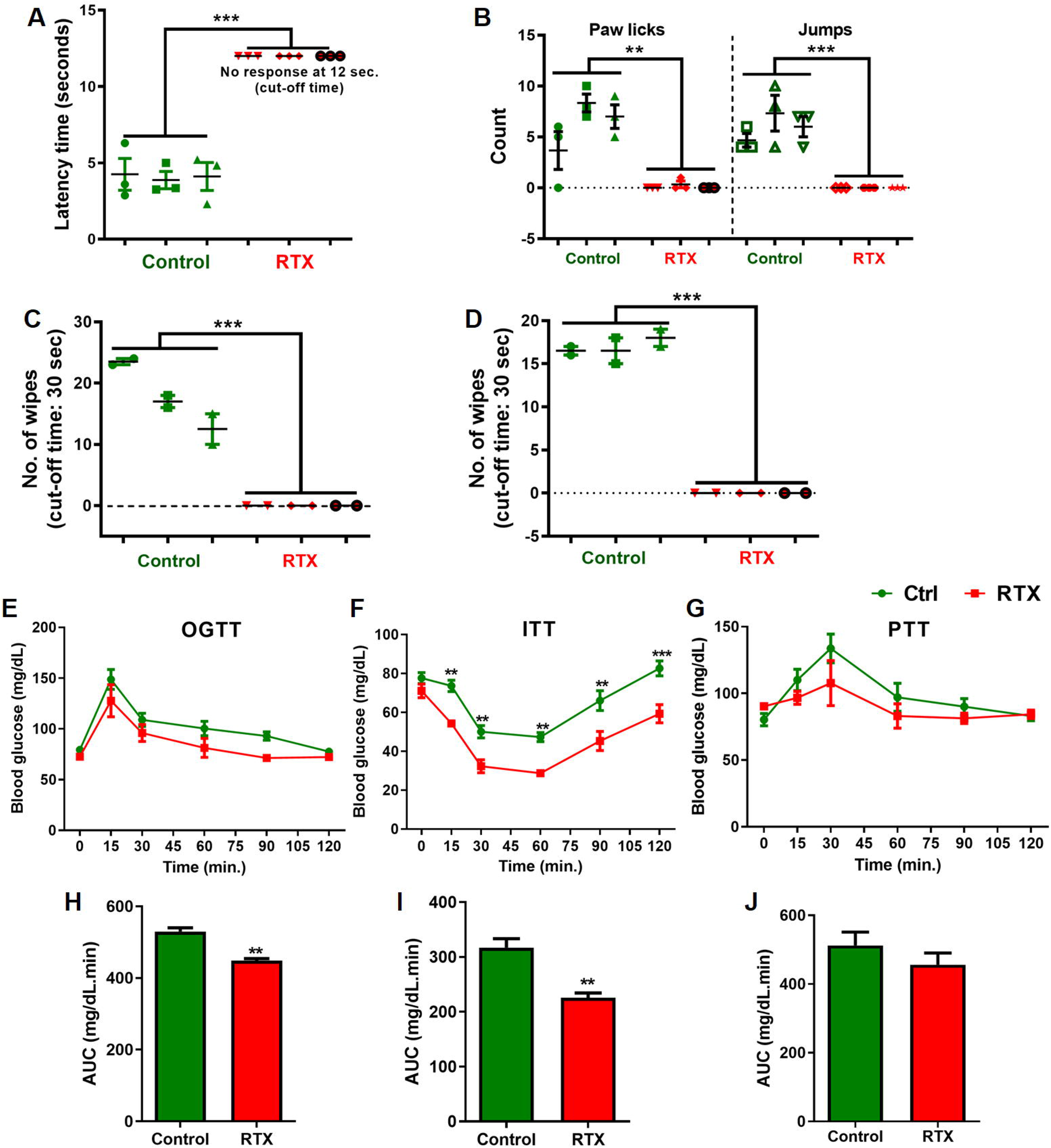
A-D) Behavioral pain assays for validating TRPV1 denervation in RTX-treated rats and E-J) glucose homeostasis tests (OGTT, PTT, ITT). All animals were fed normal pellet diet for 4 weeks. At day 0, treatment group animals received s.c. injection of 300μg/kg RTX. Physiological tests for denervation were performed 24 hours after injection, and eye wipe test was performed before sacrifice too. A) Tail flick test (2-3A current, cut-off time=12 sec., 3 reps at 5 min. intervals); B) Hot plate test (temp.=55±5°, cut-ff time=30 sec.); C) Eye wipe test (0.02%w/v capsaicin, cut-off time=30 sec.); D) Final eye wipe test in 4^th^ week (before sacrifice). Oral glucose tolerance test (OGTT), Insulin tolerance test (ITT) and Pyruvate tolerance test (PTT) were done during 3^rd^ and 4^th^ weeks of study. Animals were fasted for 12h (OGTT, PTT) or 6h (ITT). Doses: 2g/kg body weight p.o. glucose, 1U/kg i.p. insulin and 1g/kg i.p. sodium pyruvate; blood glucose levels were measured at 0 (before treatment), 15 min, 30 min, 60 min, 90 min and 120min after treatment. E-F) Recorded glucose levels and H-J) AUC analyses of OGTT, ITT and PTT respectively. Values are expressed as individual readings with mean bars. Intergroup variations were assessed by student’s t-test with Welch’s correction. Statistical significance is represented as ** P < 0.01, *** P < 0.001.

OGTT, ITT and PTT were performed to examine the effects of TRPV1 chemo-denervation on glucose homeostasis. The elevation in blood glucose levels in response to oral glucose or intraperitoneal pyruvate administration was decreased in RTX group animals (Figures 2E and 2G). Besides, insulin-induced drop in blood glucose was significantly higher in RTX group (Figure 2F), indicating hypersensitivity to insulin. AUC analyses of OGTT, ITT and PTT (Figure 2H-2J) showed an overall decrease in blood glucose levels in treatment group, suggesting changes in glucose homeostasis patterns. These findings were in agreement with a recent study conducted in mice, where capsaicin-induced TRPV1 denervation produced same effects (21). Hence, selective denervation of TRPV1^+^ neurons by RTX induced decrease in blood glucose levels.

### 3.2. RTX treatment led to changes in gene expression patterns in DRGs

To examine the effect of RTX on different classes of neurons in DRGs and colon tissue (submucosa), qPCR was performed for selected marker genes. The heatmap in Figure 3A depicts an overview of gene expression patterns of all the marker genes tested in mentioned samples. Among the selected markers for TRPV1-expressing peptidergic neurons, *Tlr4*, *Asic3*, *Ntrk1* and *Tac1* were significantly downregulated by RTX treatment. Also, in non-peptidergic neuron markers, significant decrease in *Tgr7*, *P2rx3* and *Gfra2* genes in RTX group was observed (Figure 3B). There were changes in markers for neurofilament cells too, however non-significant. In the colon samples, all the peptidergic markers showed patterns similar to those in DRGs, with Spp1 showing significant reduction. Besides, neurofilament marker *Ldhb* showed increased expression in RTX group (Figure 3C). These results indicate that while RTX-induced chemo-denervation caused loss of TRPV1-expressing peptidergic neurons, other genes were also affected, in both DRGs and colon.

**Figure 3.**
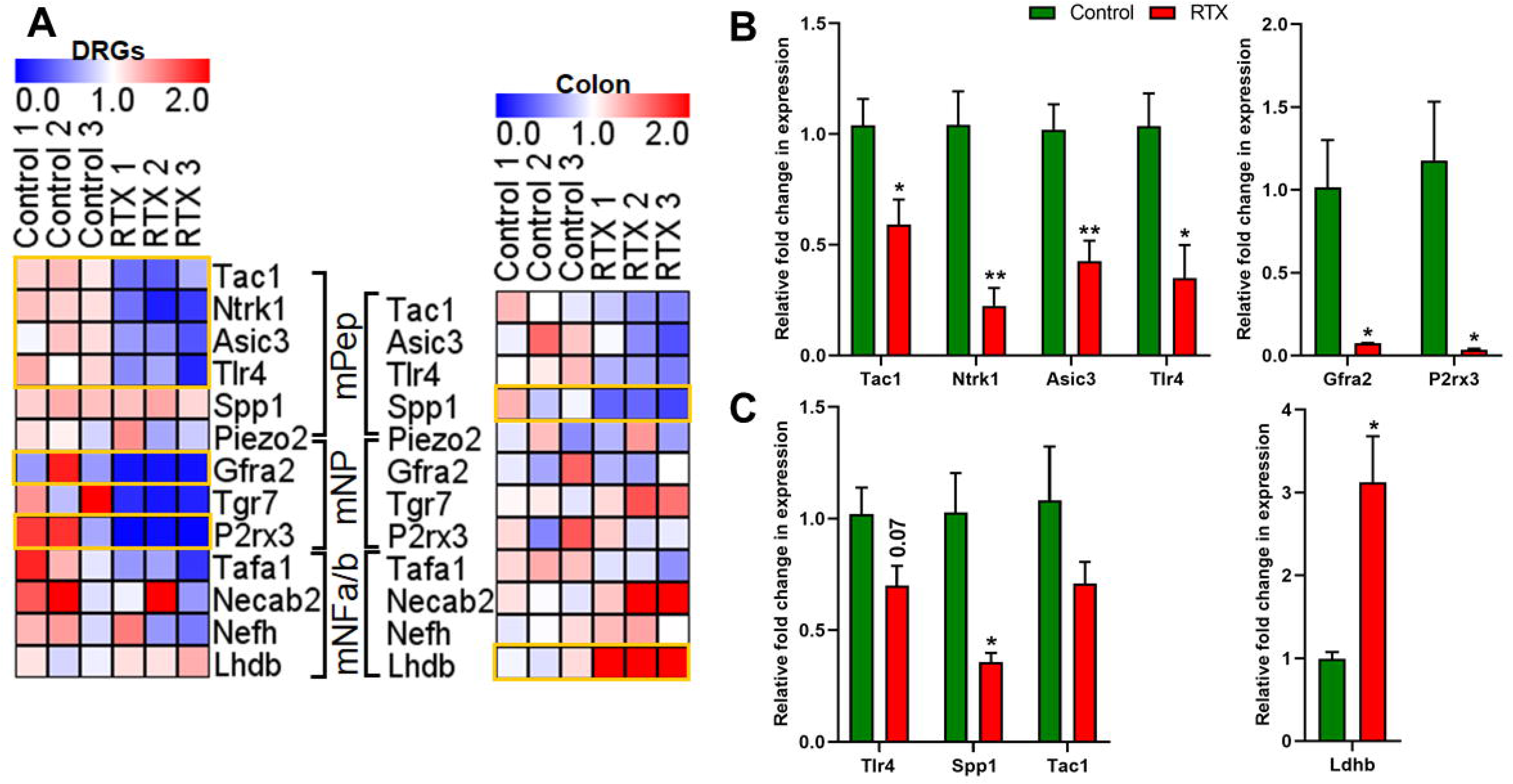
Gene expression analysis in DRGs and colon samples. All animals were fed normal pellet diet for 4 weeks. At day 0, RTX group animals received s.c. injection of 300μg/kg resiniferatoxin. After sacrifice, tissues were harvested and RNA was isolated manually using Phenol-Chloroform-Isopropanol method. After cDNA synthesis, qPCR was performed for various genes. Data was analyzed using ΔΔCt method given by Livak *et al* (2001). A) Heatmaps showing expression overview of genes in DRGs (left) and colon (right) samples, under mentioned categories – mPep: peptidergic neuron markers, mNP: non-peptidergic neuron markers, mNFa/b: neurofilaments a-type/b-type markers (genes with major changes in expression highlighted in yellow boxes); B) Genes with major changes in expression in DRGs samples; C) Genes with major changes in expression in colon samples. Values are expressed as Mean±SEM. Intergroup variations were assessed by student’s t-test with Welch’s correction. Statistical significance is represented as *P<0.05, ** P < 0.01.

### 3.3. TRPV1 ablation inhibited mucogenesis in colon

Histological analysis of colon and ileum sections was done to examine the morphological changes in gut tissues. It revealed severely decreased mucus levels in goblet cells, evident by visibly less Alcian blue uptake in RTX group colon samples (Figure 4A). Alcian blue stain intensity analysis in multiple sections using ImageJ, also showed significantly less staining in colon samples from RTX group, however these changes were not seen in ileum sections (Figure 4B).

**Figure 4.**
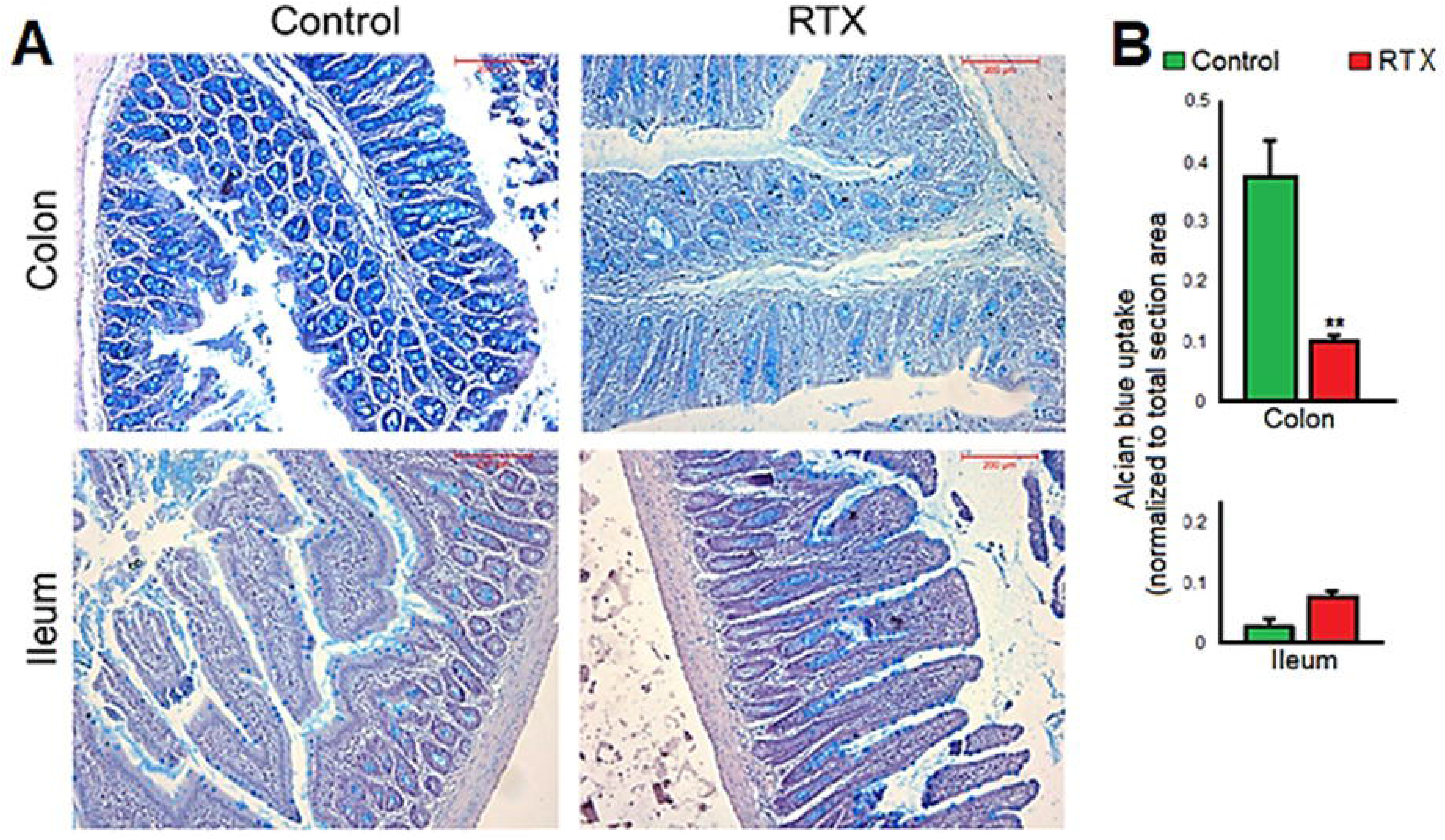
Histological analysis of mucin in colon and ileum samples. All animals were fed normal pellet diet for 4 weeks. At day 0, RTX group animals received s.c. injection of 300μg/kg resiniferatoxin. After sacrifice, tissues were fixed, paraffin-embedded and stained with Alcian blue-Hematoxylin-Eosin combination. The images were analyzed using ImageJ software (n=7-9 each). A) Representative images of colon and ileum sections from rats without (Control) or with (RTX) resiniferatoxin treatment; B) Alcian blue intensity analysis (normalized to whole section area), indicating relative amounts of mucus (glycoproteins) in the tissue. Values are expressed as Mean±SEM. Intergroup variations were assessed by student’s t-test with Welch’s correction. Statistical significance is represented as ** P < 0.01.

Gene expression analysis was performed using qPCR to investigate the changes in goblet cell marker genes, mucin genes and glycosylation enzymes genes in colon and ileum samples (Figure 5A). Various genes from these categories showed decreased expression in RTX treatment. The goblet cell marker genes – *Cdx2, Dll4, Foxa2, Klf1*; predominant mucin gene in gut – *Muc2*; genes for various mucin glycosylation enzymes – *Galnt1, Galnt2, Galnt3, Galnt5, Galnt6, Galnt7*, were significantly downregulated (Figure 5B). However, the changes in gene expression in ileum samples were not found prominent. The gene expression analysis was in accord with the histological findings. These findings indicate the role of TRPV1 in regulation of mucus secretion in gut, as evident by the diminished mucus production and downregulation of multiple associated genes in animals with TRPV1 ablation.

**Figure 5.**
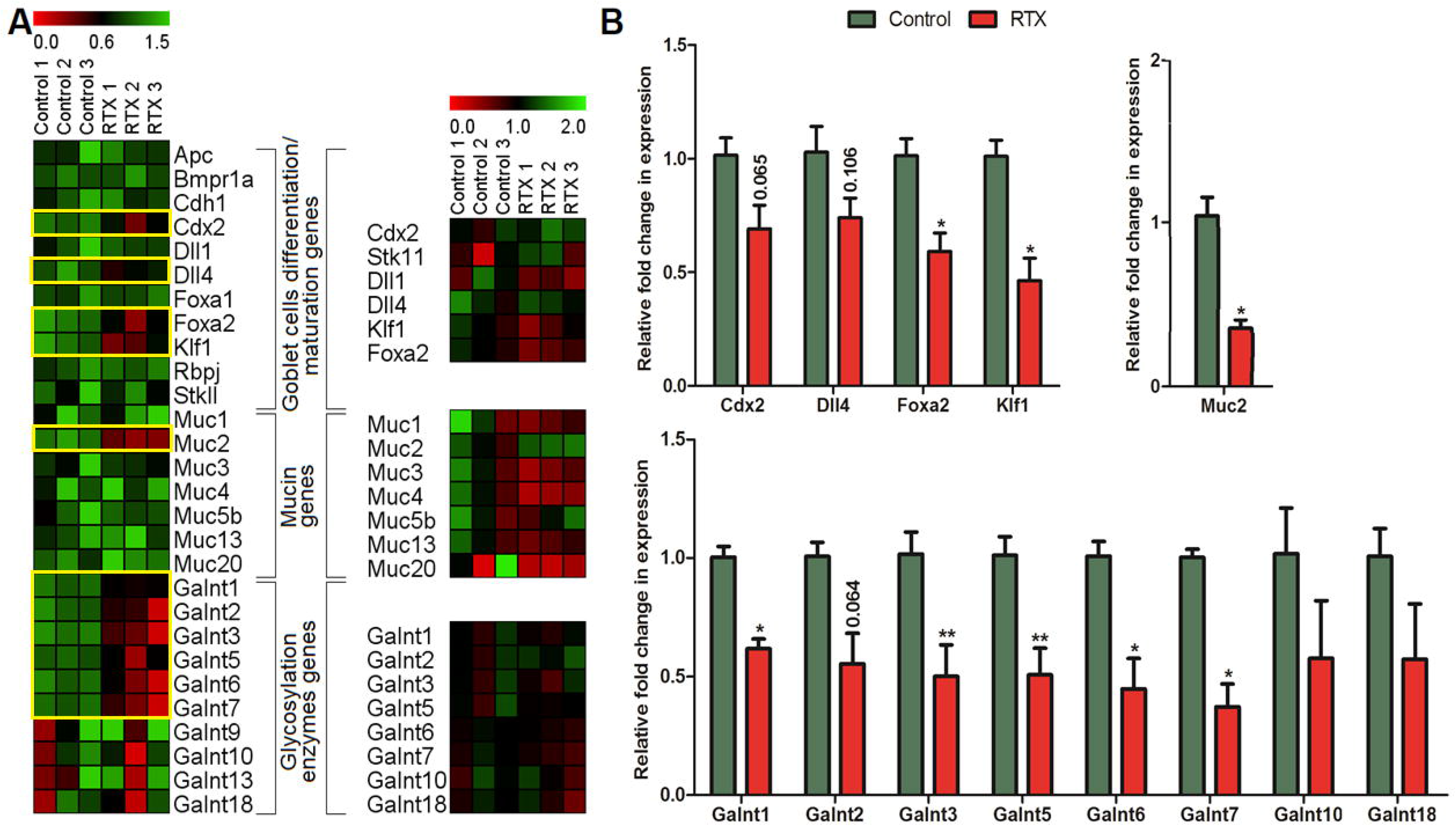
Gene expression analysis in colon and ileum samples. All animals were fed normal pellet diet for 4 weeks. At day 0, RTX group animals received s.c. injection of 300μg/kg resiniferatoxin. After sacrifice, tissues were harvested and RNA was isolated manually using Phenol-Chloroform-Isopropanol method. After cDNA synthesis, qPCR was performed for various genes. Data was analyzed using ΔΔCt method given by Livak *et al* (2001). A) Heatmaps showing expression overview of genes in colon (left) and ileum (right) samples, under mentioned categories (genes with major changes in expression highlighted in yellow boxes); B) Genes with major changes in expression in colon samples. Values are expressed as Mean±SEM. Intergroup variations were assessed by student’s t-test with Welch’s correction. Statistical significance is represented as *P<0.05, ** P < 0.01.

### 3.4. TRPV1 denervation caused decrease in bacterial metabolites and affected bacterial abundance

The concentrations of major bacterial metabolites in gut – acetate, propionate and butyrate were estimated in cecal content of rats using HPLC. There was a significant reduction in acetate and butyrate levels in RTX group (Figure 6A), which led us to further investigate the gut microbiota profile of the samples. 16S metagenome analysis was done and abundances of various bacteria were found to be significantly different in RTX group. At phylum level, the major phyla – Bacteroidetes and Firmicutes were checked. RTX group showed an increase in Bacteroidetes and decrease in Firmicutes (Figure 6B). However, the changes were not found significant. Various beneficial butyrate-producing genera from the class Clostridia – Clostridium sensu stricto, Clostridium XIVa, Clostridium XIVb, Clostridium XVII, showed major decrease in abundance in RTX group (Figure 6D), corroborating with SCFAs profiling in HPLC. Another beneficial genus – Lactobacillus, along with its various species *L. intestinalis, L. reuteri, L. gasseri*, was depleted in RTX group (Figure 6E). Besides, other genera like Anaerobacter, Lachnospiracea_incertae_sedis, Lactovum, Mucispirillum, Murimonas, Paracubacteria_genera_incertae_sedis, Peptococcus, Sporobacter showed significantly altered abundances in RTX group, compared to control (Figure 6C). These results suggest major changes in the gut microbiota profile of rats due to RTX-induced chemo-denervation.

**Figure 6.**
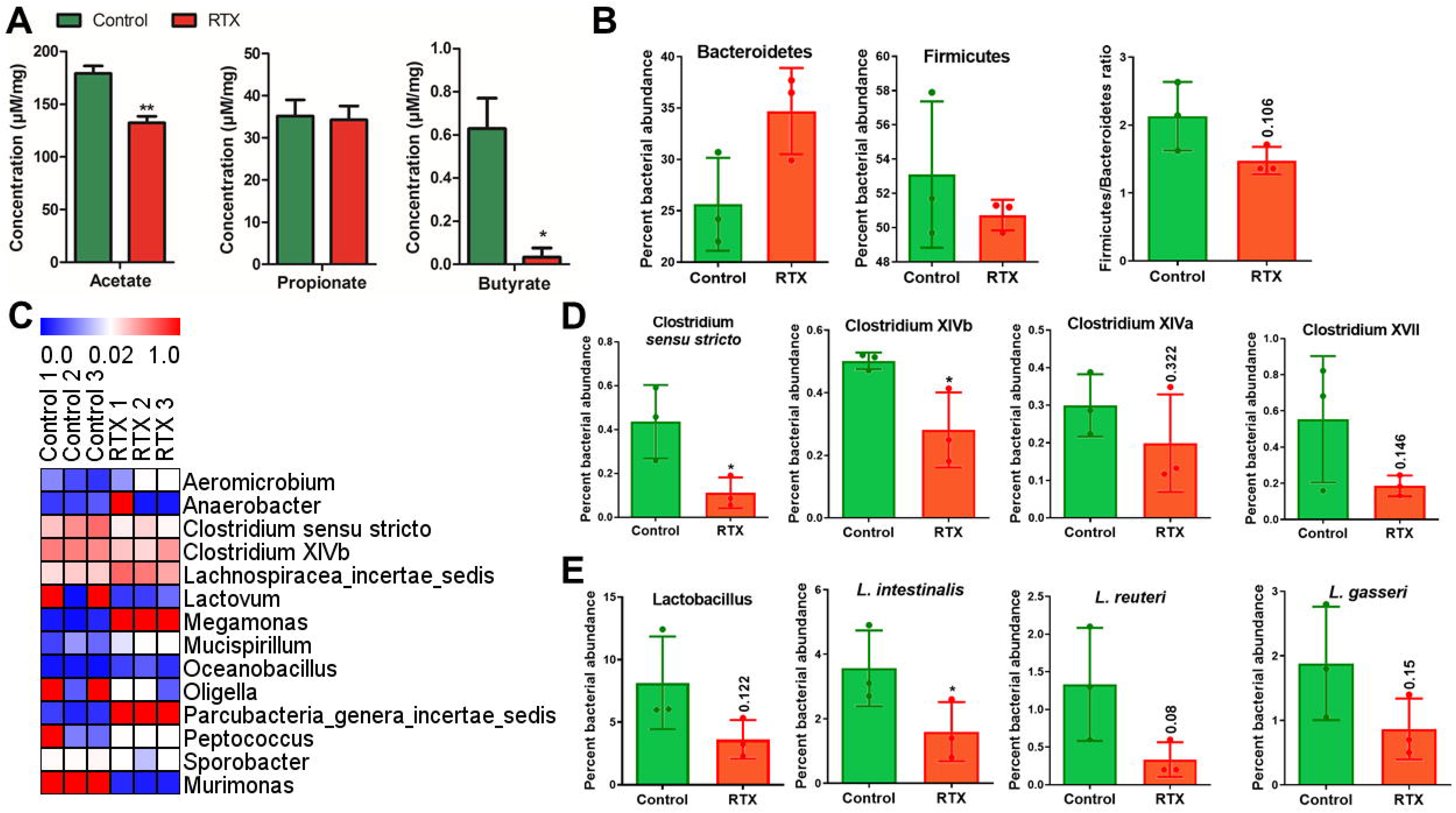
Bacterial metabolites (SCFAs) and 16S metagenome analysis in cecal content. All animals were fed normal pellet diet for 4 weeks. At day 0, RTX group animals received s.c. injection of 300μg/kg resiniferatoxin. After sacrifice, the cecal content was collected and subjected to HPLC analysis (for short chain fatty acids) and 16S metagenome analysis (for bacterial abundance). A) Concentrations of major bacterial metabolites (acetate, propionate, butyrate) in cecal content of Control and RTX group; B) Percent abundances and ratio of major gut bacterial phyla (Bacteroidetes and Firmicutes); C) Heat map showing percent abundances of genera with major changes in RTX group; D) Graphs showing changes in percent abundances of butyrate-producing beneficial bacteria from class Clostridia; E) Graphs showing changes in percent abundances of genus Lactobacillus, and species from same genus related to human gut. Values are expressed as Mean±SEM. Intergroup variations were assessed by student’s t-test with Welch’s correction. Statistical significance is represented as numbers (non-significant) or *P<0.05, ** P<0.01.

## 4. DISCUSSION

Since the discovery and understanding of its function, TRPV1 has been extensively studied for its therapeutic and preventive actions against multiple diseases, primarily as analgesics and anti-cancer drugs (Bujak et al., 2019; Premkumar and Sikand, 2008; Szallasi et al., 2007). After showing early promise, development of TRPV1 antagonists as analgesics was abruptly halted due to serious side effects associated to it, *i.e.* hyperthermia. The expansive physiological involvement of TRPV1 makes it difficult to succeed in such strategies without side-effects (Premkumar and Sikand, 2008; Szolcsanyi and Sandor, 2012). On the other hand, TRPV1 modulation by dietary components such as capsaicin (Szallasi, 2005; Yang and Zheng, 2017), piperine (Szallasi, 2005), gingerol (Yin et al., 2019) etc gave a new direction to TRPV1 oriented research. Capsaicin has been studied a lot in relation to inflammation and pain relief, and is also being investigated for its role in metabolism and related disorders like obesity (Baskaran et al., 2019; Fattori et al., 2016; Narang et al., 2017).

TRPV1 is reported to be expressed primarily in the sensory neurons innervating most of the organs, including gastrointestinal tract (Holzer, 2011). Studies have shown that oral administration of capsaicin induced beneficial effects in obesity, such as increase in energy expenditure and browning in white adipose tissues, decreased lipid accumulation, and also improvement in gut dysbiosis conditions and changes in bacterial metabolites composition (Baboota et al., 2014; Shen et al., 2017; Wang et al., 2020). One of the important findings was increase in abundance of *Akkermansia muciniphila*, a mucin-feeding bacterium, which has been shown to improve overall microbial profile in gut (Baboota et al., 2014; Shen et al., 2017). As mucus layer deterioration has been associated with high-fat diet induced gut derangements (Birchenough et al., 2017; Everard et al., 2013), and the literature also shows a connection between capsaicin administration, TRPV1 and increase in expression of major mucins like MUC2, MUC5AC (Kang et al., 1995; Yang et al., 2013), we hypothesized that the effects of capsaicin in gut might be TRPV1-mediated and related to improved mucus secretion..

The previously reported effects of TRPV1 denervation – loss of nociception towards capsaicin and heat (Almasi et al., 2003; Bates et al., 2010) were clearly observed, suggesting that the denervation did remove the TRPV1^+^ cells of treated animals. Also, we found major decrease in overall blood glucose and significantly increased insulin sensitivity in RTX-treated rats, which has also been reported in a recent study as an effect of loss of TRPV1 by chemo-denervation (Bou Karam et al., 2018). Further, we also studied the gene expression pattern in DRGs connected to the sensory afferent nerves present in colon (Hockley et al., 2019). It has been shown that gene markers of peptidergic (*Tac1, Ntrk1, Asci3, Tlr4*) and non-peptidergic *(Gfra2, Tgr7, P2rx3)* class of sensory neurons were significantly decreased whereas there was no significant change in gene markers of neurofilament types a and b *(Tafa1, Necab2, Nefh, Lhdb)*. Overall, these confirmations suggested ablation of TRPV1+ neurons.

The change in mucin secretion and related phenomenon has a reported effect on gut microbiota. Alternations such as decrease in mucin layer can induce gut dysbiosis and *vice versa* (Corfield, 2018; Sicard et al., 2017). Upon histological examination using Alcian blue, we found that there was a drastic decrease in mucus production in colon tissues of RTX-treated animals. To explore it at gene expression level, we performed qPCR for numerous genes involved in goblet cell maturation, mucin expression and glycosylation of mucins. Major genes involved in goblet cell maturation, like *Cdx2, Dll4*, *Foxa2, Klf1* (Katz et al., 2002; Pellegrinet et al., 2011; Yamamoto et al., 2003; Ye and Kaestner, 2009), were downregulated in RTX group. Most mucin genes showed no significant change, but *Muc2*–the predominant mucin gene in gut (Tytgat et al., 1994), showed significantly decreased expression. Gal-NAc is the most abundantly present in O-type glycans associated with mucins (Brockhausen et al., 2009) Therefore, we examined the expression of various genes for glycosylation of mucins with particularly Gal-NAc. Most of these genes were found downregulated, *Galnt2, Galnt3, Galnt5, Galnt6, Galnt7* showing significant decrease in expression. This further elucidated our findings in histological examinations. These results show that there was a major impairment in the mucus production mechanism in gut, particularly at glycosylation steps, due to TRPV1 denervation, which indicates the involvement of TRPV1 in maintenance of mucus production.

Decrease in mucin secretion is related to altered gut microbiota profile. To understand how neuronal ablation of TRPV1 affected the gut microbial population, we performed 16S metagenome analysis in cecal content of animals. We found major changes in bacterial profile – including significant changes in abundance of genera like Anaerobacter, Lachnospiracea_incertae_sedis, Lactovum, Mucispirillum, Paracubacteria_genera_*incertae_sedis*, Peptococcus, Sporobacter and Murimonas. Further, RTX caused significant decrease in genera like Clostridium *sensu stricto*, Clostridium XIVa, Clostridium XIVb and Clostridium XVII, which have major role in butyrate production (Pryde et al., 2002; Van den Abbeele et al., 2013). Butyrate is a bacterial metabolite known to have nourishing effects on various metabolic parameters, including promotion of gut mucus secretion (Bedford and Gong, 2018). Analysis also showed decreased overall abundance of beneficial lactic acid bacteria, including *L. intestinalis, L. reuteri* and *L. gasseri*, reflecting a compromised gut health in RTX group. *Lactobacillus spp* are reported to promote mucus secretion via producing metabolites like oxo-fatty acids (Kim et al., 2017), and also acts as probiotic and imparts gut health benefits (Kechagia et al., 2013). In another interesting finding, the Firmicutes/Bacteroidetes ratio in RTX group was decreased, which is reported to have opposite pattern in disorders like obesity (Castaner et al., 2018). Also, there was decrease in the levels of SCFAs in RTX treated animals, particularly butyrate, which consolidated our findings from metagenomic analysis. Overall, we can argue that TRPV1 ablation resulted in compromised mucin production mechanisms along with gut dysbiosis and decrease in healthy bacterial metabolites. An interesting take on these findings, as suggested in a recent study is that TRPV1 denervation by chemical means, while causes a systemic removal of TRPV1^+^ neurons, does not affect its expression in the non-neuronal cells (Kun et al., 2012). Considering this information, it seems that while TRPV1 may not be directly involved in the mentioned effects *via* ion transport, the activation of TRPV1^+^ nerve cells and subsequent release of various neurotransmitters might have more to do with these changes. Multiple studies have shown that these neurotransmitters have modulatory effects on mucus secretion patterns through localized action in gut. In DRGs samples of RTX-treated animals, we found decreased expression of peptidergic neuron markers like *Tac1, Ntrk1, Asic3, Tlr4* and non-peptidergic markers *Gfra2, P2rx3*. Also, in RTX-colon samples, decreased expression of *Tac1, Spp1, Tlr4 etc* was observed. *Tac1* encodes for precursors of various tachykinins, including Neurokinin 1 and Substance P (SP). Both of these have been reported to stimulate mucus discharge from airway goblet cells (Chu et al., 2000; Kuo et al., 1990). SP plays crucial role in inflammatory and immune responses, mainly *via* vasodilation control (O’Connor et al., 2004). While SP has not been particularly associated with mucus secretion in gut, evident decrease in its neuronal as well as non-neuronal expression suggests a correlation. *Asic3*, a gene coding for major class of acid sensing ion channels, is also implicated in protective function, including maintenance of bicarbonate and mucus secretion (Holzer, 2015). *Tlr4* is vital in immune response, it mediates the dissolution of bacterial lipopolysachharides and consolidates gut barrier (Fukata and Abreu, 2007). Similarly, *Gfra2* and *P2rx3* have been associated with immune response and neurotransmitters release (Coutinho-Silva et al., 2005; Rossi et al., 2003). *Spp1* codes for osteopontin, which is also involved in mucosal protection and inflammatory response (Chen et al., 2013). *Ldhb*, a neuofilament marker and gene for enzyme involved in glycolysis, showed increased expression in RTX-treated colon samples, reflecting on results from blood glucose tests. It has also been implicated in tumor progression. To sum up, our findings show that by targeting TRPV1^+^ neurons through desensitization or denervation, we risk to impair multiple physiological mechanisms.

TRPV1 desensitization or blocking has been and is still proposed as preventive or therapeutic option for treatment of various diseases. A recent study suggested the development of TRPV1 antagonists as anti-diabetics, based on the TRPV1 ablation-induced decrease in glucose levels (Bou Karam et al., 2018), as also observed in our experiment. But simultaneously, the derogatory changes occurred in gut environment, clearly seen in our study, indicate that great care needs to be taken in such advances, emphasizing on site-specific and more precisely targeted approaches towards TRPV1 modulation.

In conclusion, we report for the first time that systemic TRPV1 chemo-denervation leads to severe impairment in gut mucus production and causes gut dysbiosis, which indicates that TRPV1 is essentially involved in these phenomena. We propose that interaction of TRPV1 and the products of its modulation with non-neuronal cells (like goblet cells) and gut microbiota might be the key to better understanding the role of TRPV1 in these physiological processes. Finally, we provide evidence to be considered, when developing TRPV1 antagonism/desensitization strategies for different diseases, to limit their side effect profile.

## ACKNOWLEDGMENT

Authors would like to thank Department of Biotechnology, Government of India for research grant given to National Agri-Food Biotechnology Institute (NABI) and Dr. Mahendra Bishnoi. Authors would like to thank University Grant Commission (UGC) Government of India for research fellowship given to Mr. Vijay Kumar. Authors would like to thank Mr. Rakesh Maurya, NxGenBio Life Sciences, 120, 3rd floor, Hargovind Enclave, New Delhi-110092, India for carrying out metagenome analysis.

## AUTHOR’S CONTRIBUTION

VK and MB designed experiments, VK, PK and NM carried out experiments, VK and MB did analysis of experiments, PK and KKK reviewed and edited the paper, VK and MB wrote and edited the paper, MB Lead contact

## CONFLICT OF INTEREST

Authors declare no competing financial interest

## Notes

### Competing Interest Statement

The authors have declared no competing interest.

